# The power and limitations of species tree-aware phylogenetics

**DOI:** 10.1101/2023.03.17.533068

**Authors:** Tom A. Williams, Adrian A. Davin, Benoit Morel, Lénárd L. Szánthó, Anja Spang, Alexandros Stamatakis, Philip Hugenholtz, Gergely J. Szöllősi

## Abstract

Species tree-aware phylogenetic methods model how gene trees are generated along the species tree by a series of evolutionary events, including the duplication, transfer and loss of genes. Over the past ten years these methods have emerged as a powerful tool for inferring and rooting gene and species trees, inferring ancestral gene repertoires, and studying the processes of gene and genome evolution. However, these methods are complex and can be more difficult to use than traditional phylogenetic approaches. Method development is rapid, and it can be difficult to decide between approaches and interpret results. Here, we review ALE and GeneRax, two popular packages for reconciling gene and species trees, explaining how they work, how results can be interpreted, and providing a tutorial for practical analysis. It was recently suggested that reconciliation-based estimates of duplication and transfer frequencies are unreliable. We evaluate this criticism and find that, provided parameters are estimated from the data rather than being fixed based on prior assumptions, reconciliation-based inferences are in good agreement with the literature, recovering variation in gene duplication and transfer frequencies across lineages consistent with the known biology of studied clades. For example, published datasets support the view that transfers greatly outnumber duplications in most prokaryotic lineages. We conclude by discussing some limitations of current models and prospects for future progress.

**Significance statement:** Evolutionary trees provide a framework for understanding the history of life and organising biodiversity. In this review, we discuss some recent progress on statistical methods that allow us to combine information from many different genes within the framework of an overarching phylogenetic species tree. We review the advantages and uses of these methods and discuss case studies where they have been used to resolve deep branches within the tree of life. We conclude with the limitations of current methods and suggest how they might be overcome in the future.

## Introduction

Species tree-aware methods based on probabilistic gene tree-species tree reconciliation have recently emerged as a powerful approach in phylogenomics and comparative genomics. Recent studies have used reconciliation methods, including the tools ALE (Szöllõsi et al. 2013), GeneRax (Morel et al. 2020) and SpeciesRax (Morel et al. 2022), to infer the root of species trees (Williams et al. 2017; Coleman et al. 2021; Cerón-Romero et al. 2022), map the evolutionary origins of gene families (David & Alm 2011; Martijn et al. 2020; Schön et al. 2022), estimate more accurate single gene trees and ancestral sequence reconstructions (Groussin et al. 2015; Blanquart et al. 2021), and draw conclusions about the contributions of gene gain (Dharamshi et al. 2023), transfer, duplication and loss to the evolution of bacterial, archaeal (Sheridan et al. 2020, 2022) and eukaryotic (Szöllősi et al. 2015; Harris et al. 2022) genomes.

Compared to traditional approaches such as concatenation, species tree-aware methods based on probabilistic reconciliation models such as ALE have a number of advantages for inferring species trees. In particular, their ability to account for gene origination, duplication and horizontal transfer allows more of the genome to be included in analyses. This is particularly useful in the context of microbial evolution, where often only a small proportion of genes evolve vertically (Doolittle 1999; Lerat et al. 2005; Dagan & Martin 2006, 2007; Treangen & Rocha 2011; Tria & Martin 2021), and are therefore amenable to concatenation. By estimating and comparing probabilities for different scenarios of gene duplication, transfer and loss events under any root, these approaches allow species trees to be rooted without the use of an outgroup. These methods have therefore been applied to rooting questions when no obvious outgroup is available or when the only available outgroup stems from a different domain of life (Williams et al. 2017; Coleman et al. 2021), as outgroups that are only distantly related to ingroup taxa can lead to mis-rooting due to long-branch attraction (Bergsten 2005; Kapli et al. 2021; Williams et al. 2021). In addition, methods such as ALE and GeneRax infer events of duplication, transfer and loss directly from the data (gene trees or multiple sequence alignments), without the need for prior assumptions about their rates. Other methods, such as TreeFix-DTL and ecceTERA, that infer parsimony-based reconciliations (Doyon et al. 2011; Bansal et al. 2018; Jacox et al. 2016), require the costs of such events to be specified prior to analysis.

Recently, some of us applied the ALE reconciliation approach to root the phylogeny of Bacteria (Coleman et al. 2021). By using a model that accounts for transfers, duplications and losses, we were able to use a substantially greater amount of the available data (11,272 bacterial gene families, in comparison to the <60 vertically-evolving genes that can be used to infer the unrooted tree of life (Harris et al. 2003; Gribaldo et al. 2010; Spang et al. 2015; Hug et al. 2016; Parks et al. 2020; Martinez-Gutierrez & Aylward 2021; Moody et al. 2022)) to investigate the position of the root. These analyses supported a basal divergence between two major bacterial lineages (clans), the Gracilicutes (Gibbons & Murray 1978) and the Terrabacteria (Battistuzzi et al. 2004; Battistuzzi & Hedges 2009), consistent with other recently published species trees (Raymann et al. 2015; Martinez-Gutierrez & Aylward 2021; Moody et al. 2022; Taib et al. 2020; Aouad et al. 2022). Mapping traits onto these rooted phylogenies represents one approach to understanding the nature of the last bacterial common ancestor, which can be compared to alternative approaches that polarise evolution by identifying major transitions (Cavalier-Smith 2006) or that do not rely on a rooted species tree (Xavier et al. 2021).

However, the use of ALE in this and other analyses has recently been criticised. Tria and Martin (2021) criticised the rates of gene duplication and transfer (and in particular, the ratios of these rates) inferred using ALE because they were inconsistent with the large excess of gene transfers over duplications in prokaryotic genomes frequently observed in previous analyses (Lerat et al. 2005; Treangen & Rocha 2011; Tria & Martin 2021). In a subsequent paper (Bremer et al. 2022), these and other authors argued that transfer and duplication rate ratios in ALE analyses were unrealistic, and that these biases affect the inference of rooted species trees using the ALE model.

To address these criticisms, we first describe the reconciliation model underlying ALE (Szöllõsi et al. 2013) and GeneRax (Morel et al. 2020), explain how it works and how the results can be interpreted. Based on this understanding, we summarise what published analyses have concluded about variation in the processes of molecular evolution across the tree of life. We also clarify a number of potential misconceptions about these methods and their results in the recent critiques (Tria & Martin 2021; Bremer et al. 2022), and show that reconciliation-based inferences about frequencies of duplication, transfer and loss are in good agreement with previous results using other methods. Finally, we provide a practical guide for researchers who wish to use reconciliation tools to perform phylogenomic and comparative genomic analyses, review current limitations and suggest future directions for addressing them.

## Results and Discussion

### A primer on gene tree-species tree reconciliation

*Species trees* describe the history of ancestor-descendant relationships that relate modern organisms to the root of the tree. The internal nodes of species trees (common ancestors of extant taxa) are of great interest to evolutionary biologists because they correspond to ancestral species that no longer exist. Unless fossil data are available, the only information we have to study these ancestors and infer their characteristics derives from their living descendants, in particular their genome sequences (Zuckerkandl & Pauling 1965; Boussau & Daubin 2010).

*Gene trees* describe the evolutionary history of individual gene families that trace their history back to a single common ancestor. Internal nodes in gene trees represent the divergence of gene lineages that often accompany species-level divergences but can also represent gene duplications or the coalescence of distinct alleles in a population.

In some cases, particularly for essential and rarely-transferred gene families such as ribosomal proteins, gene trees can be adequately modelled as following the species tree of the organisms that harbour them, and gene evolution follows along the species tree. Most often, however, gene and species trees differ, for at least three reasons. First, gene trees might disagree with each other and the species tree due to errors, weak or insufficient signal and uncertainties in phylogenetic reconstruction: trees are statistical inferences and are not guaranteed to be correctly or fully resolved. Second, speciation can fail to perfectly sort genetic variation from the parent population into the daughter species when several speciation events happen in quick succession or population sizes are very large. If recombination is sufficiently strong, the ancestral population may contain a large number of independent gene lineages. As a result, ancestral genetic variation may persist in descendant lineages and the divergence of some genes will pre-date that of the species in which they reside today, potentially exhibiting alternative evolutionary relationships. This phenomenon is called incomplete lineage sorting; for example, about 30% of the human genome is more closely related to gorilla (Scally et al. 2012) than to chimpanzee, and 0.5% is more closely related to orangutan than to either chimpanzee or gorilla, due to incomplete lineage sorting (Hobolth et al. 2011). Finally, gene and species trees can differ due to processes such as gene duplication, gene loss, and gene transfer — acquisition of genes from a source other than a direct ancestor, for example through active uptake of environmental DNA or through infection by a virus or genetic element that may carry genetic material from an organism it infected previously (reviewed in (Soucy et al. 2015)). The spread of antibiotic resistance among Bacteria is perhaps the most prominent example of gene transfer, but it occurs extensively across the tree of life, particularly in prokaryotes, and affects most or all classes of genes ((Gogarten & Townsend 2005; Soucy et al. 2015; Daubin & Szöllősi 2016; Irwin et al. 2021)). Lineage-level processes such as hybridization (Moran et al. 2021) and endosymbiosis (Timmis et al. 2004; Nelson-Sathi et al. 2012; Martin et al. 2015) induce large-scale gene transfer and are major causes of gene tree-species tree discordance.

Traditionally, disagreement between gene and species trees has been viewed as presenting a serious challenge to our ability to infer meaningful species trees at any level, including the universal tree of life (Doolittle 1999; Creevey et al. 2004; Dagan & Martin 2006). However, such disagreements and more generally discordant evolutionary histories across genes and genomes, can also be viewed as one of the most useful sources of information about evolutionary history. For example, the occurrence, extent and timing of interbreeding between humans, Neanderthals and Denisovans has been investigated by comparing genetic variation among modern human populations and ancient genomes (reviewed in (Racimo et al. 2015). More broadly, the logic of gene tree-species tree reconciliation — that is, the idea that it is misleading to view some gene trees as agreeing or disagreeing with the species tree, because all gene trees have evolved along the same species tree (Maddison 1997) - has a long history in phylogenetics. In many cases, key events in the history of a gene family can be discerned by informally interpreting the gene tree in the context of prior knowledge about species-level relationships. For example, the statistically supported nesting of aphids within fungi in gene trees for carotenoid biosynthesis (Moran & Jarvik 2010) supported the hypothesis that aphids acquired these pigments from fungi by HGT because the alternative explanation - ancestral presence of carotenoids in the opisthokont ancestor of animals and fungi, followed by widespread and repeated loss in most lineages since - seems implausible. HGT across larger evolutionary distances is easier to detect from disagreements between gene and species trees: for example, carotenoid biosynthesis in a disparate group of protists (single-celled eukaryotes) also appears to result from HGT from prokaryotes to eukaryotes and then within eukaryotes (Rius et al. 2023). The grouping of nucleotide transport proteins used to steal host ATP from bacterial and fungal intracellular parasites in single gene trees supported the hypothesis of HGT from bacteria to fungi (microsporidians), facilitating adaptation of the fungi to the intracellular niche (Dean et al. 2018). Similarly, the acquisition of bacterial genes for aerobic metabolism by haloarchaea was supported by the nesting of archaeal sequences within the Bacteria in single gene trees (Nelson-Sathi et al. 2012; López-García et al. 2015; Martijn et al. 2020).

In addition to identifying HGTs, gene tree-species tree discord can also help to interpret patterns of gene loss and duplication. The presence of the genetic toolkit to produce stomata (gas exchange pores) in mosses, combined with a plant species phylogeny rooted between bryophytes and tracheophytes, provided evidence that stomata were already present in the earliest land plants, but subsequently lost in some descendant lineages (Chater et al. 2016; Harris et al. 2020; Clark et al. 2022). At the deepest level of the tree of life, the inference that some genes duplicated prior to the divergence of Archaea and Bacteria provided a means to root the entire tree, because each paralogues (duplicated gene copy) can act as an outgroup to root the species-level relationships reflected in the other (Iwabe et al. 1989; Gogarten et al. 1989; Brown & Doolittle 1995; Zhaxybayeva et al. 2005; Dagan et al. 2010).

Models for gene tree-species tree reconciliation take this same logic — using explicit evolutionary events (gene origins, duplications, transfers and losses) to explain gene tree-species tree discord — and apply it systematically to large, genome-scale datasets in an automated way. The main strengths of these approaches are that they are objective and scalable, in the sense that they can be applied systematically to many gene families to pool evolutionary signal. There are many ways to explain the evolution of a family of homologous genes given a species tree, and the most probable gene tree, together with the most likely reconciliation is usually unclear. Basing inference on one gene tree, or one optimal reconciliation, often involves making arbitrary choices between statistically indistinguishable alternatives. It is therefore useful to treat the problem statistically, explicitly quantifying the uncertainty of estimates at each level and, as explained below, inferring reconciliations using a model of the evolutionary process in which the probabilities of each type of event can be learned from the data.

In practical terms, the main limitations of current reconciliation-based inference methods are of two kinds. First, in the interest of computational tractability, current models make simplifying assumptions that are often violated by real data. For example, the reconciliation model used by ALE and GeneRax (described below) includes only one duplication (delta, ς), transfer (τ, tau), and loss (λ, lambda) parameter for each gene family (that is, a total of 3 parameters describing the relative branch-wise probabilities of duplication, transfer and loss for each family; see below for more detail on how the model is parameterised). In reality, DTL frequencies vary across clades. For example, vertically inherited bacterial endosymbionts and endoparasites often experience reduced selection and are characterised by high gene loss (McCutcheon & Moran 2011). The impact of these simplifying assumptions is not well understood, but - as in other branches of phylogenetics - it seems likely that the accuracy of inferences will improve as more sophisticated models that better fit the data are developed (Kapli et al. 2021; Williams et al. 2021). ALE and GeneRax also do not model incomplete lineage sorting (ILS), which may be a frequent source of gene tree-species tree conflict in recombining (e.g., sexual) lineages.

The second limitation of current approaches is that, as with any complex bioinformatics pipeline, there is the potential for the introduction of error at each step. For instance, methods for inferring gene families (clustering and functional annotation approaches) are far from perfect. More generally, complex pipelines that chain together multiple tools are susceptible to bugs in individual tools, which can be difficult to detect. Readers who have inferred single gene trees will be aware that gene families vary greatly in levels of conservation (both sequence and functional), evolutionary mode and phylogenetic informativeness, and at present it is difficult to identify clustering settings that work for all (or most) families. Indeed, this is a general limitation of current high-throughput comparative genomic analyses, and an area where progress is badly needed. In practice, one approach might be to experiment with different procedures and compare results. Moreover, the impact on the final results of varying the plethora of *ad hoc* parameters and default threshold settings in the core bioinformatics tools of complex data analysis pipelines is rarely assessed. Ideally, one should generate an ensemble of plausible datasets for downstream analysis by varying these parameters — for instance, as in the Muscle 5 MSA tool that returns an ensemble of plausible alignments (Edgar 2022) or the Bayesian method bali-phy that samples both alignments and gene trees using MCMC (Redelings 2021).

### Inference of gene duplication, transfer and loss events using ALE and GeneRax

We begin by explaining how gene duplication, transfer and loss events are inferred using the probabilistic reconciliation method used in both ALE and GeneRax. A full treatment of the model, including the likelihood function that is optimised during inference, is provided in (Morel et al. 2020), while additional algorithmic details of how ALE calculates the likelihood can be found in (Szöllõsi et al. 2013).

ALE and GeneRax implement a probabilistic model that explains how a gene family can evolve inside a species tree. The process being modelled is one in which a gene family appears on a given branch of the species tree and then evolves from there following events of vertical descent, gene transfer, duplication and loss. The probabilities with which these events occur are estimated from the data. If a time-calibrated species tree is available (for example, has been inferred using a molecular clock analysis), then reconciliation can be performed in a time-consistent manner; that is, gene transfers into the past are not permitted. If a dated species tree is unavailable or not known with confidence, reconciliation can be performed using “undated” algorithms (e.g., ALEml_undated, “undated ALE” below) that do not make assumptions about the temporal order of speciations. In undated algorithms, transfers into direct ancestors are not permitted but are otherwise not guaranteed to be time-consistent because the method does not know the relative age of nodes that are not in a direct ancestor-to-descendant line on the rooted species tree. Published analyses of microbial data have generally used undated algorithms due to the difficulty of inferring reliable time trees for microbes.

In the case of ALE, the gene trees to be reconciled have been previously computed. Normally, the user will use a distribution of these gene trees (such as the one that can be obtained from bootstrap or the posterior chain of a Bayesian analysis) that captures the uncertainty in the tree topology from the alignment. ALE proceeds to “decompose” the splits that have been found in the distribution of gene trees and reconcile that distribution with the species tree. The resulting reconciliation is a new gene tree that can potentially include splits found in different trees from the original distribution. In the case of GeneRax, the inference of the gene tree is simultaneous with the reconciliation. When new topologies are proposed, the likelihood search accounts for both the probability of the tree topology given the alignment and the probability of that tree topology given the species tree.

When performing these reconciliations, ALE and GeneRax map the gene tree into the species tree “backwards” (from the tips of the gene tree — which can be unambiguously mapped to the tips of the species tree — to the root of the gene tree). As the algorithm works back from the tips, DTL events are exhaustively enumerated. Since the algorithm is probabilistic, different reconciliations can occur with different probabilities. As a result of the exhaustive enumeration of reconciliations, ALE can be used to efficiently sample reconciliations after DTL parameter optimization. While ALE outputs by default 100 reconciliations, the user can change this to a larger value with minimal computational overhead. The origination point of the gene family is a part of each particular reconciliation, and sampling reconciliations can result in a distribution of potential origination points on the branches of the species tree. In fact, ALE and GeneRax account for both the uncertainty in the reconciliation and the uncertainty in gene tree inference (that is: given a multiple sequence alignment, various gene trees are possible; given each of those gene trees, various reconciliation scenarios are possible). Inferences about the origination point of the gene and the inferred number of DTL events that occur are averaged over the sampled reconciliations to explicitly quantify uncertainty. For example, consider a gene family in which half of the sampled reconciliations involved a gene transfer, while half did not; this family would have 0.5 inferred transfers.

Next, we describe how the model is parameterised. The key events of the undated ALE/GeneRax DTL model are gene duplication (D), transfer (T) loss (L) and speciation (S) with the root node of the gene tree corresponding to an origination event in the species tree; speciation refers to vertical descent from an ancestral node to its immediate descendant. The probabilities with which a gene present on a branch on the species tree will experience each of these events is described by the three parameters ς, τ, and λ. In particular, the probabilities of duplication (*p*^D^), transfer (*p*^T^), loss (*p*^L^) or vertical descent (i.e. speciation given by *p*^S^) on a branch can be written:

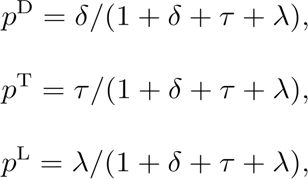

and

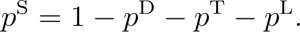

As outlined above, the process is that a gene family originates at some internal branch of the species tree, then experiences events according to the above discrete state stochastic process on each subsequent branch, before one or more copies arrive at the tips of the species tree (Figure 1). Note that ς, τ, and λ parametrize the relative probability of vertical descent versus D, T, or L on each species tree branch, so they cannot be interpreted as rates (numbers of events that occur in per unit time), and are not directly proportional to the number of inferred events. The numbers of D, T, and L events for each gene family are inferred by averaging the events that occur over the distinct reconciliation scenarios (ways that the gene tree can be drawn within the species tree using a series of gene birth and death/DTL events) that are sampled according to their probability given information on the gene tree and the values of ς, τ, and λ. The parameters of the model are optimised by jointly maximising (i) the phylogenetic likelihood on the multiple sequence alignment and (ii) the likelihood of possible reconciliation scenarios. These two components of the joint likelihood - the phylogenetic likelihood and the reconciliation likelihood - can be thought of as expressing two different aspects of the uncertainty of the reconciliation, given a multiple sequence alignment and a species tree. The phylogenetic likelihood captures the relationship between the multiple sequence alignment (MSA) and the gene tree: a given MSA might have evolved along many different gene trees, each with a different probability. The reconciliation likelihood captures the relationship between the gene tree and the species tree: given a gene tree and a species tree, there are many different ways to draw the gene tree into the species tree (many different possible reconciliations), each with a different probability. When performing the reconciliation, we need to account for both of these sources of uncertainty: the uncertainty in the gene tree, given the MSA; and the uncertainty in the reconciliation, given the gene tree.

**Figure 1:**
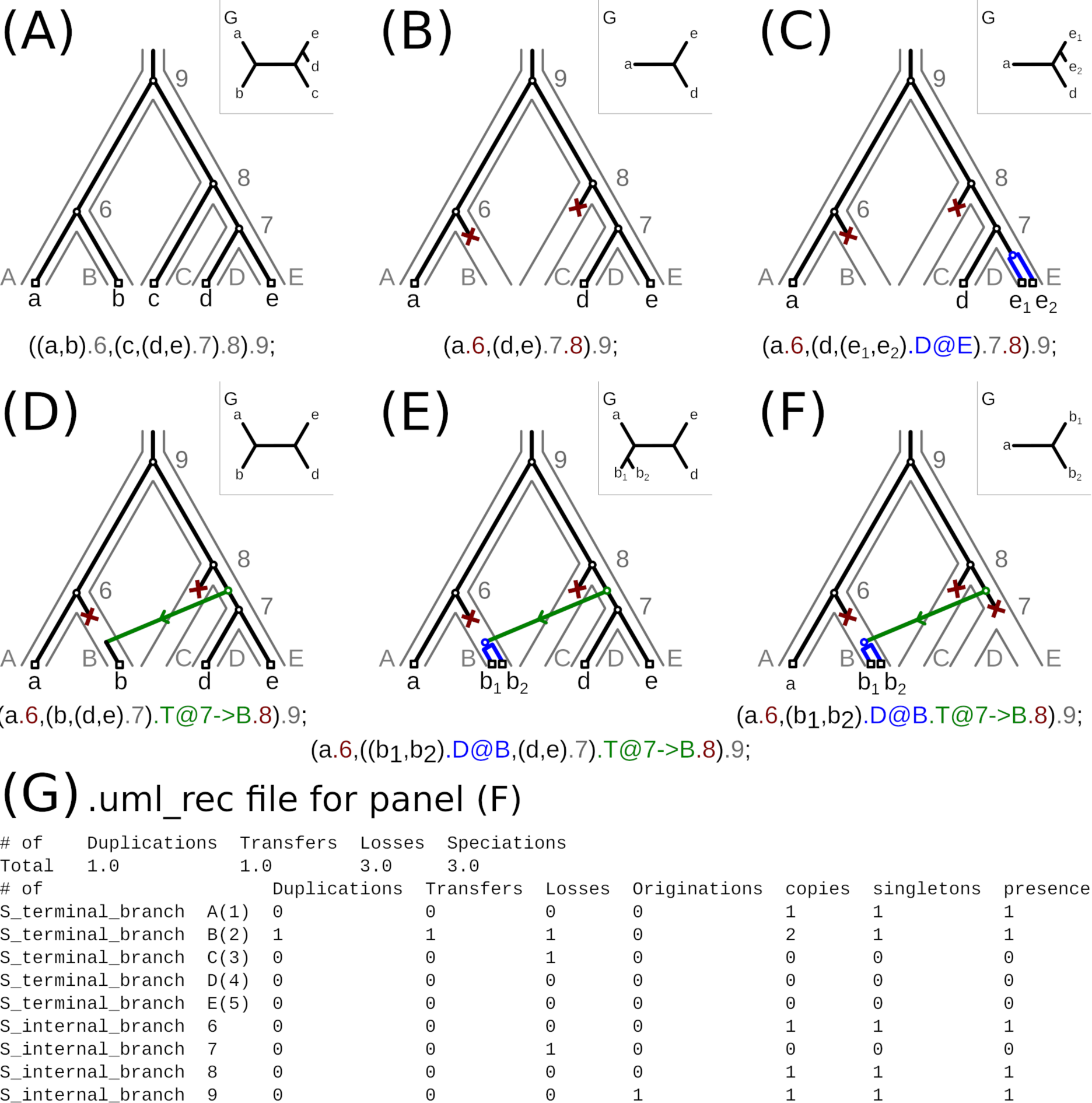
Interpreting ALE output. Possible reconciliations of different gene trees given a species tree and the extended Newick string representations for duplication, transfer, loss and speciation events. The species tree’s topology with node names (leaf names and node numbers) is depicted in grey, the gene tree in black (also depicted separately for each case in the top right corner). Evolutionary events needed to reconcile the gene and species trees are highlighted in different colours: red for gene loss, blue for gene duplication, green for gene transfer and a black circle for speciation. Terminal nodes (leaves or tips) are drawn as black squares. (A) the gene tree topology is congruent with the species tree, so no evolutionary events are required to reconcile them. (B) the gene tree does not include sequences from species B and C, which can be explained by speciation and loss (SL) events on the species tree. (C) depicts a gene duplication (D event) on the branch leading to E. (D) shows a transfer (T event) from branch number 7 to the terminal branch B. (E) shows a transfer from branch 7 to branch B and duplication on branch B (DT event). (F) depicts all three events at once: a transfer followed by a loss on branch 7 and a duplication on the receiver branch B abbreviated as DTL event. Finally, panel (G) shows the output file *.uml_rec generated by ALEml_undated for the gene tree-species tree reconciliation depicted in panel (F). The uml_rec file contains a summary of the observed evolutionary events, in the case of panel (F) 1 duplication, 1 transfer, 3 losses and 3 speciations. After this, a list of Newick strings for each sampled reconciled gene tree follows, in the format shown beneath panels (A)-(F). The uml_rec file ends with a description of the frequency of observed events per branch and with other branch-wise statistics: branch category, branch name or numeric ID, duplications, transfers, losses, originations, copies, singletons and presence. These events can be summarised (for example, summed per-branch over all gene families) to compute the total number of events of each type on a branch. We provide scripts to tabulate these summaries in the accompanying Github repository (https://github.com/AADavin/ALEtutorial).

ALE and GeneRax treat the calculation of the joint likelihood differently: GeneRax calculates this joint (phylogenetic and reconciliation) likelihood directly from the multiple sequence alignments of each gene family, whereas ALE approximates the phylogenetic likelihood using conditional clade probabilities computed from a distribution of gene family trees. In both cases, explicit evolutionary scenarios involving a series of gene birth and death events that have given rise to the genes in extant genomes (that is, *reconciliations*) can be sampled according to their probability. These reconciliations can then be summarised to extract information about the inferred number of gene duplication, transfer and loss events that occurred during the history of the gene family, and their mapping onto the rooted species tree (see Figure 1).

The approximation used in ALE to sample the distribution of gene trees has both advantages and disadvantages. The main disadvantage is that an additional step is required in the analysis (either an MCMC search or the inference of a bootstrap distribution), and accurately computing these distributions can be challenging, especially on large numbers of taxa where the number of tips in the single gene trees can be large compared to the length of the multiple sequence alignment. The degree of difficulty for a phylogenetic inference on a given MSA or essentially lack of signal for obtaining a stable single gene (family) tree can now be predicted via machine learning methods (Haag et al. 2022). However, because the approximation needs only to be computed once for each gene family, ALE is faster than GeneRax for the purpose of evaluating different rooted species trees (or root positions on a single unrooted topology). By using an approximation based on a distribution of gene trees that allows the efficient exploration of the space of gene trees (Szöllõsi et al. 2013), ALE also sums over all gene tree topologies and roots. In contrast, GeneRax conditions on the ML rooted gene tree, and therefore accounts for the uncertainty in single gene tree topologies to a lesser degree. Further, because the input gene tree distributions can be generated modularly using any tool, ALE also allows the use of parameter-rich site-heterogeneous substitution models (such as the CXX (Quang et al. 2008), EDM (Schrempf et al. 2020), PMSF (Wang et al. 2018) or MAMMAL (Susko et al. 2018) models or the CAT model implemented in PhyloBayes (Lartillot & Philippe 2004) that often fit real data better than the simpler standard alternatives. ALE might therefore be particularly useful in studies of ancient evolutionary relationships, where substitution model fit is known to be important (Williams et al. 2021; Kapli et al. 2021). By contrast, GeneRax is faster than ALE for analyses on a single rooted species tree, making it ideal in cases where the species tree is known and the primary aim of the analysis is to infer accurate gene family trees. Simulations, results on real data (Szöllõsi et al. 2013; Scornavacca et al. 2015; Morel et al. 2020), and empirically assayed biochemical properties of ancestrally reconstructed proteins based on alternative gene trees (Groussin et al. 2015) suggest that, by making use of the additional information from the species tree, reconciliations methods including GeneRax and ALE infer more accurate single gene trees than approaches based on the phylogenetic likelihood alone.

In both ALE and GeneRax, the gene tree-species tree reconciliation is performed on a fixed rooted species tree. To test different root positions, the analysis must be run once for each candidate rooted species tree; the gene family likelihoods obtained with each root can then be compared using a tree selection test (such as the Approximately-Unbiased (AU) test (Shimodaira 2002)) to identify a confidence set of roots. For example, Coleman et al. (2021) evaluated support for 62 root positions on an inferred unrooted tree of Bacteria, and could reject a root position on all but three adjacent branches (a “root region”) that had the three highest summed gene family log-likelihoods; a step-by-step guide to this procedure was described in a recent book chapter (Harris et al. 2022); see also the ALE tutorial (https://github.com/AADavin/ALEtutorial) that accompanies this manuscript. Note however, that exhaustively evaluating root positions in this way can be compute-intensive for large datasets.

### Alternatives to reconciliation for rooting phylogenetic trees

Given the biological interest of rooting problems, many alternative approaches to outgroup rooting are being developed. One class of methods makes use of branch length information to root trees. Building on the idea of midpoint rooting (rooting a tree in the middle of the longest tip-to-tip path), MAD (Tria et al. 2017) and MinVAR (Mai et al. 2017) are methods that root trees at the position that implies the minimum variation in molecular evolutionary rate from the root to the tips. Molecular clock models (Ho & Duchêne 2014; dos Reis et al. 2016) can also use branch length information to root trees, although in practice these models are not often used for rooting, but rather to infer divergence times on a fixed, rooted species tree. A second class of rooting methods makes use of asymmetric or non-reversible features of the substitution process. For example, the NONREV (Naser-Khdour et al.) and UNREST (Yang 1994) models, implemented in IQ-TREE 2 (Minh et al. 2020) and RootDigger (Bettisworth & Stamatakis 2020), relax the assumption of reversibility in the standard GTR substitution model, so that the instantaneous rate of change from, say, A to G is different to that from G to A. As a result, the likelihood of observing the multiple sequence alignment given the tree also depends on the root of the tree, allowing the root to be inferred without assuming an outgroup. While the best outgroup-free rooting approach is debated and may be dataset-dependent, previous work suggests that all of these approaches can capture root signal and correctly root trees under some conditions (Bettisworth & Stamatakis 2020; Tria et al. 2017; Coleman et al. 2021; Wade et al. 2020; Naser-Khdour et al.; Dombrowski et al. 2020).

To evaluate the extent of agreement between different outgroup-free rooting approaches on an interesting test dataset, we applied MAD and the non-reversible NONREV+G model to the bacterial dataset we analyzed previously using ALE (Coleman et al. 2021). As shown in Figure 2, root support is significantly correlated between ALE, MAD and NONREV+G, with all three approaches favouring a similar set of root positions. We compared the values from the three methods using the Spearman’s rank correlation, finding a value different from zero in the three cases (MAD vs NONREV ρ = −0.27, *p* = 0.03; MAD vs ALE ρ = −0.48, *p* = 5.56 · 10^−5^; NONREV vs ALE: ρ = 0.71, *p* = 1.03 · 10^−10^ Of the two probabilistic methods, ALE has greater power to reject root positions with lower log-likelihoods using an Appoximately-Unbiased (AU) test (Shimodaira 2002) (Figure 2). This might reflect the difference in the nature of root signal being captured by these two approaches: the summed ALE log-likelihood pools root signal from reconciliations across a large number of gene families (11,272 gene families in this case), while the NONREV+G log-likelihoods summarise the information about the non-reversibility of the substitution process in the 62-gene concatenated alignment. Overall, the degree of agreement observed is particularly encouraging given that the three methods make use of largely distinct sources of root information, and suggests that analyses combining different types of root information are a promising direction for future progress. For example, ALE could be used to reconcile gene tree distributions rooted using MAD, NONREV or UNREST.

**Figure 2:**
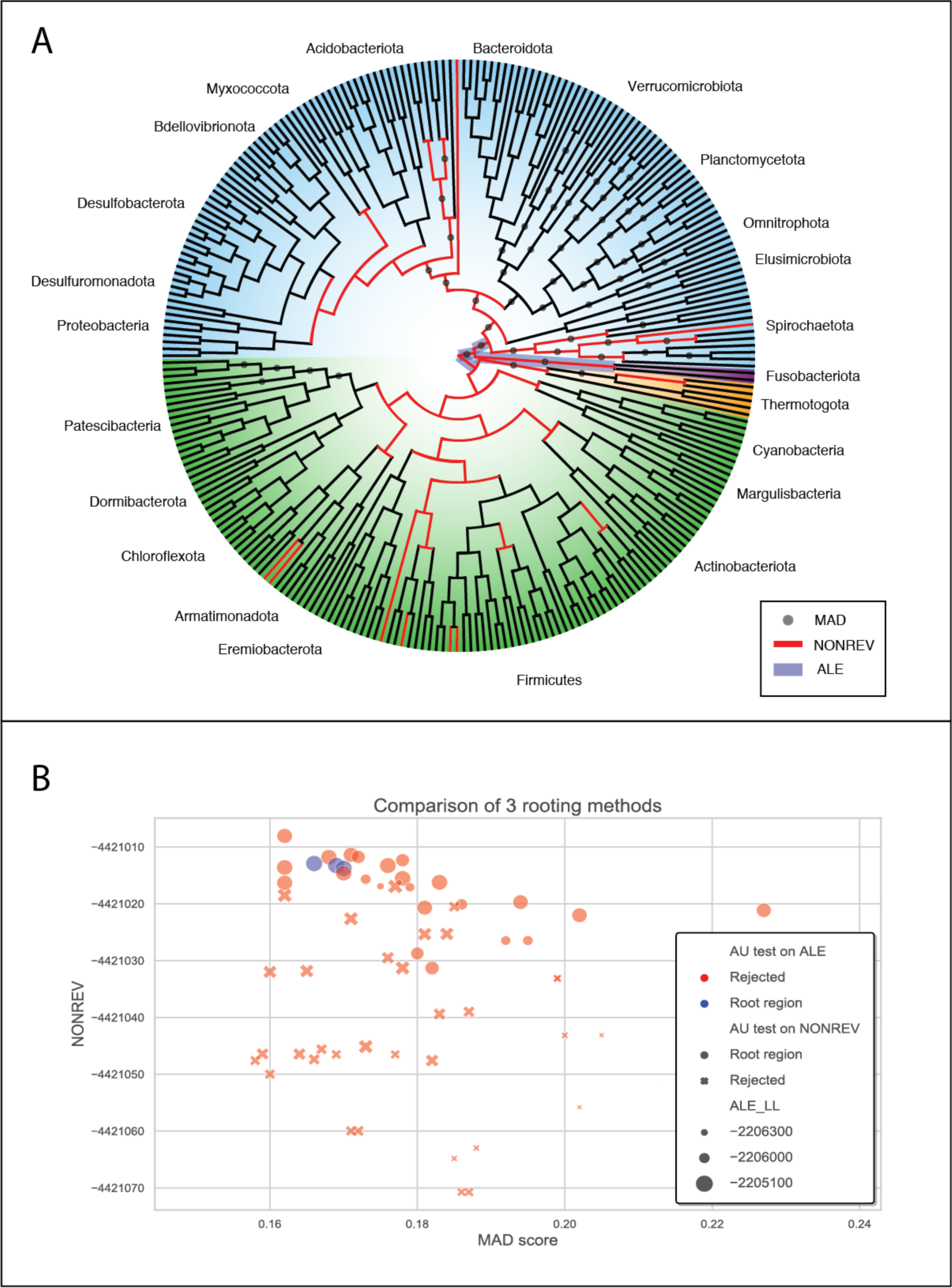
Agreement between reconciliations, branch lengths, and a non-reversible substitution model on the position of the bacterial root. (a) An unrooted cladogram of Bacteria indicating root support from ALE, MAD and NONREV+G. Terrabacteria are highlighted in green, Gracilicutes in blue. For the likelihood-based methods, root positions that could not be rejected by an AU-test (p < 0.05) are indicated. An AU test using ALE log-likelihoods rejected all but three of the internal branches as a plausible root position, whereas NONREV+G log-likelihoods were more equivocal. This might be because the ALE analysis makes use of more data (11,272 gene families compared to a 62-gene concatenation). For the MAD analysis, we plot the nodes with the 10% lowest (best) AD scores. (b) Agreement between MAD scores, ALE reconciliation log-likelihoods, and NONREV+G log-likelihoods for the internal nodes of the bacterial species tree; scores from the three methods are significantly correlated (see main text).

### Accuracy of reconciliation-based analyses on simulated and real data

Method accuracy can be investigated using simulations or by analysis of empirical datasets for which there is general agreement on the most plausible answer. Both approaches have limitations: in simulation experiments, the correct answer (in this case parameter values, numbers of events, inferred tree or root position) is known, and so accuracy can be quantified unambiguously. However, simulated data do not fully capture the complex patterns and biases in real data that can cause methods to fail, and similarity between the models used to simulate and analyse the data can give false confidence in the accuracy of an approach as the line of argument is, in a sense, circular. Empirical data are of course more realistic, but performance is difficult to benchmark objectively because the truth is not known.

In previous studies we have extensively used simulations to assess the accuracy with which ALE recovers duplication, transfer and loss events (Coleman et al. 2021; Morel et al. 2020) and the accuracy of root inference on the basis of gene family likelihoods (Williams et al. 2017). To investigate robustness to model violations, these simulations used models that were somewhat more complex than the inference model (for example, analysing the data using simpler substitution models than those used to generate the input MSAs (Szöllõsi et al. 2013; Groussin et al. 2015) or using the continuous-time Zombi simulator (Davín et al. 2020)). These analyses suggested that simulated events are recovered accurately across a range of rates, and that the reconciliation model can correctly distinguish between gene transfers and duplications, even in the presence of gene loss (Coleman et al. 2021). Analysis of simulated gene trees suggested that GeneRax and ALE could accurately infer the topology and root of single gene trees, supporting the expectation that including the species tree significantly improves accuracy (Morel et al. 2020; Coleman et al. 2021). Simulations and empirical data also suggest that ALE can reproducibly recover the species tree root using gene family likelihoods (Williams et al. 2017; Bremer et al. 2022).

An example of an empirical dataset for which there is broad consensus on the most plausible phylogeny, based on several lines of evidence, is the radiation of land plants, with a root between two major lineages - the vascular plants (tracheophytes) and the morphologically simpler bryophytes - the best-supported hypothesis (Clark et al. 2022; One Thousand Plant Transcriptomes Initiative 2019; Puttick et al. 2018; Harris et al. 2020). Consistent with this consensus, ALE recovered a root region centred on the bryophyte-tracheophyte divide (Harris et al. 2022), despite enormous variation in gene loss frequencies among early plant lineages. Reconciliation-based inference of the root of Opisthokonts (Bremer et al. 2022) represents another informative empirical case study: there is general agreement that the root of the tree lies between Fungi and Metazoa (Torruella et al. 2012), and ALE recovered this root with maximal support.

In cases where there is less community consensus on the root of the tree, there has recently been some encouraging agreement between reconciliation and more traditional phylogenetic approaches. For example, in the context of bacterial phylogeny, the placement of the long-branching and genome-reduced Patescibacteria (Candidate Phyla Radiation) as sister to the Chloroflexota+Dormibacterota within the Terrabacteria has recently gained support from both standard phylogenetic (Taib et al. 2020; Martinez-Gutierrez & Aylward 2021; Moody et al. 2022) and reconciliation-based (Coleman et al. 2021) approaches. A bacterial root at, or near, a deep divide between Gracilicutes and Terrabacteria has also received support from both reconciliation and outgroup-rooted analyses (Battistuzzi & Hedges 2009; Raymann et al. 2015; Coleman et al. 2021; Aouad et al. 2022; Martinez-Gutierrez & Aylward 2021; Moody et al. 2022). A putative archaeal root between at least some DPANN clades and other Archaea has been recovered both in reconciliation (Williams et al. 2017) and more traditional analyses (Dombrowski et al. 2020; Martinez-Gutierrez & Aylward 2021). As phylogenetic methods improve and new lineages of Archaea and Bacteria are discovered, the roots of major microbial radiations will continue to be tested (Aouad et al. 2022).

### Do prior assumptions bias the estimation of model parameters in ALE?

Having described the logic of gene tree-species tree reconciliation and the ALE/GeneRax algorithm, we now address some recent critiques that constitute apparent misconceptions about how the methods work. For readers who wish to apply these methods to their own data, we have also created a Github repository containing scripts that can be used to parse ALE output files (https://github.com/AADavin/ALEtutorial).

The first issue relates to how parameter values are estimated in ALE. Tria and Martin (2021) suggested that ALE requires the input of prior ς, τ, and λ rates, while Bremer et al. (2022) claimed that parameter estimates were biased by hard-coded 1:1τ: ς priors. In fact, model parameters in ALE and GeneRax are estimated via maximum likelihood optimization without any prior assumptions. For clarity, is is worth noting that some other reconciliation tools do make use of weights for each type of event, which can be set by the user or left as defaults (for example, the parsimony method RANGER-DTL (Bansal et al. 2018)); however, all of the analyses criticised in Tria and Martin (2021) and Bremer et al. (2022) were performed using ALE, which directly estimates these values from the data.

Bremer et al. (2022) further suggested that the default equal initial values for ς and τ-that are required by the Bio++ implementation (Guéguen et al. 2013) of the standard Nelder-Mead optimization algorithm used in ALE for maximising the likelihood - have an undue influence on the optimised values, although they did not provide any evidence to support the claim. To test whether initial values influence the ML estimates, we sampled 100 gene families at random from the bacterial dataset (Coleman et al. 2021) re-analyzed by Bremer et al., and for each family we estimated the ς, τ and λ parameters 100 times from different random starting values (chosen independently and uniformly from the interval [0.01,10.] for ς, τ and λ parameters). The results (Supplementary Figure 1) show that the optimised parameter estimates are highly robust to the starting values, with median standard deviations of 8.87 · 10^−8^, 3.14 · 10^−7^ and 1.00 · 10^−7^ in ς, τ and λ parameters. This indicates that the ML optimization algorithm used is able to find the global optimum of the likelihood in terms of the DTL parameters.

### Are ALE-based estimates of duplication, transfer and loss unrealistic?

Some previous studies (Lerat et al. 2005; Treangen & Rocha 2011; Tria & Martin 2021) have suggested that HGT is very common in archaeal and bacterial genomes but less frequent in eukaryotes. HGT occurs in eukaryotes (Husnik & McCutcheon 2017; Irwin et al. 2021) but the mechanisms and frequency are debated (Martin 2017; Leger et al. 2018). Rates of HGT also appear to vary across eukaryotic clades: for example, HGT is relatively rare in animals, perhaps as a result of the germ-soma distinction (Boto 2014), but appears to be more common in single-celled eukaryotes including Fungi (Richards et al. 2011; Bruto et al. 2014) and Rhizarians (van Hooff & Eme 2023). High-quality genomes from additional eukaryotic groups will likely help to constrain frequencies of HGT more broadly in eukaryotes.

In their recent critique of ALE, (Bremer et al. 2022) suggested that estimated rates of duplication and transfer were biologically unrealistic for two datasets from different domains of life because they failed to capture the expected difference in the dynamics of genome evolution between Bacteria (Coleman et al. 2021) and eukaryotes (Bremer et al. 2022). To evaluate these claims, we summarised the ALE output from the two datasets (Figure 3). In the bacterial dataset the median branch-wise number of transfers is in fact an order of magnitude higher than that of duplications (Figure 3) in good agreement with published analyses (Lerat et al. 2005; Treangen & Rocha 2011) and consistent with the expectation that transfers are more frequent than duplications in Bacteria (Tria & Martin 2021). The pattern observed in the opisthokont dataset (Bremer et al. 2022) is quite different. Within Metazoa, ALE infers a large excess of duplications over transfers (median branchwise T/D 0.29), while inferred transfers exceed duplications in Fungi, though not to the extent observed in the bacterial dataset (median branchwise T/D 2.36). These results are consistent with the view that the germ-soma distinction likely acts as a barrier to transfer in animals (Boto 2014), while transfers are more frequent in Fungi (Richards et al. 2011; Ocaña-Pallarès et al. 2022), and more frequent still in Bacteria.

**Figure 3:**
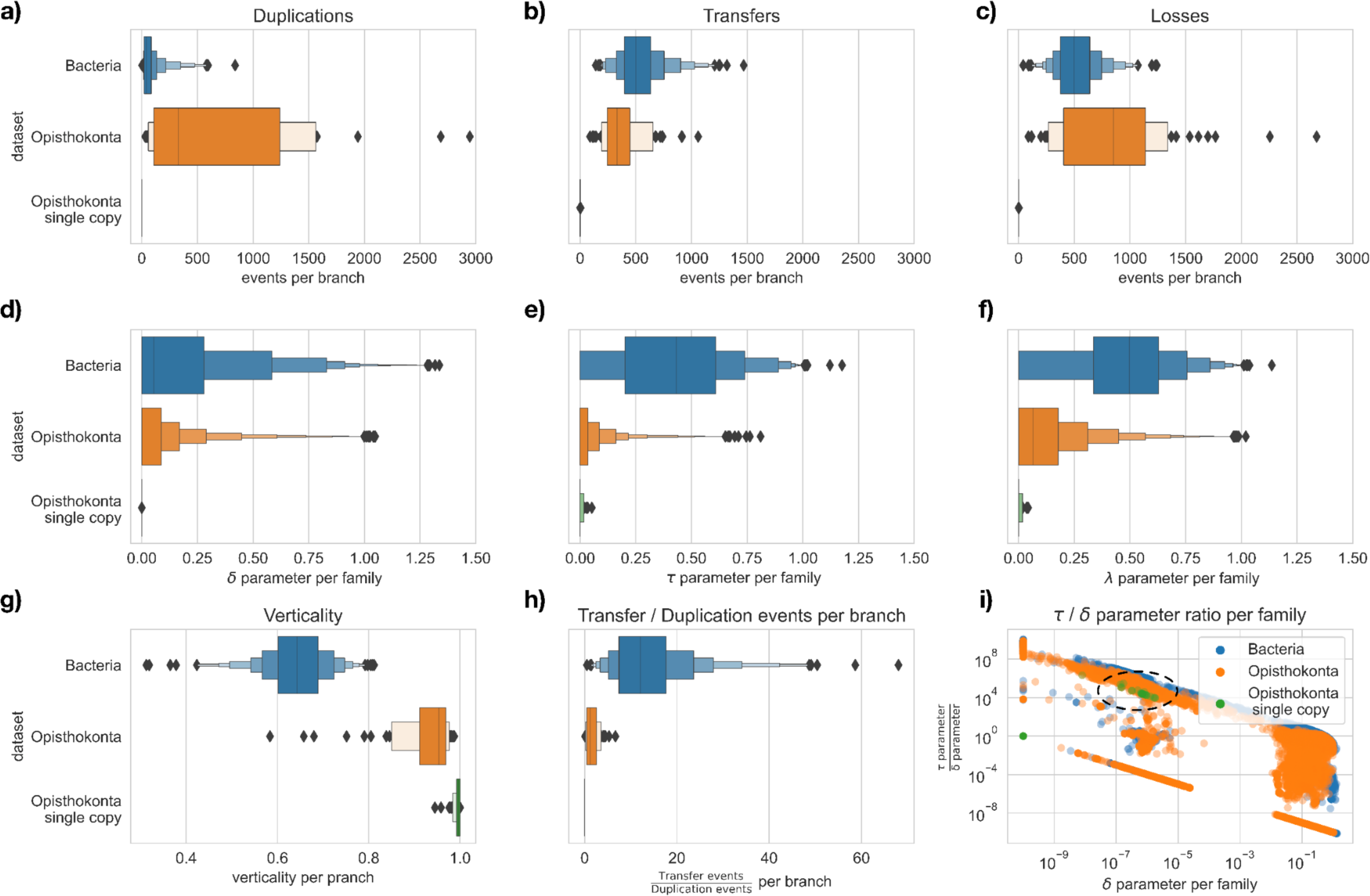
Reconciliation-based estimates of gene transfer, duplication and loss in the bacterial (Coleman et al. 2021) and opisthokont (Bremer et al. 2022) datasets. ALE reconciliation output files contain a variety of parameter values and inferences, and understanding what each represents is key to interpreting the results. (a-c) Branch-wise estimates of the number of gene duplication, transfer and loss events in the bacterial and opisthokont datasets. As expected, transfers greatly outnumber duplications in Bacteria, while the numbers of events are more balanced in the opisthokont dataset. Single-copy marker genes in opisthokonts experience 0 duplications, and indeed few transfer or loss events. (d-f) ς, and λ parameters for each gene family in the bacterial and opisthokont datasets. While genome dynamics are reflected in the distributions of per-family parameter values (for example, is generally much higher in bacteria than opisthokonts), the between-lineage patterns are less clear because the parameter distributions also reflect an enormous variation in propensity for transfer, duplication and loss across gene families. Note that parameter values cannot be interpreted as numbers of events, but describe relative probabilities within each gene family. (g) Given a species tree and a set of reconciled gene trees, branch-wise verticality can be calculated as the number of occurrences of vertical evolution from the ancestral to descendant node, divided by the sum of vertical and horizontal transfer events along the branch (Coleman et al. (2021)). Based on ALE estimates, we find that opisthokonts have much higher verticality than Bacteria, as expected (Boto 2014; Ocaña-Pallarès et al. 2022). (h) The per-branch ratio of transfer to duplication events inferred by ALE; this is a natural comparator of the per-genome counts of transfer and duplication events reported in previous analyses. As expected, T/D is higher in Bacteria than opisthokonts. Note that T/D is misleading for the opisthokont single-copy orthologous genes because no duplications were inferred in any of the 117 genes in this set. (i) The family-wise ratio of and τ and ς parameter values. This metric is highly variable, both due to biological variation in transfer and duplication frequencies across gene families (Nagies et al. 2021), but also simply because dividing by very low ς parameter values is misleading (note that τ/ς is often very high simply because ς is close to 0; see circled region in panel (i)). Note that (h) and (i) were conflated in Bremer et al. (2022), leading the authors to conclude that ALE-based ratios of transfer and duplication were unrealistic (see Supplementary Text for further discussion).

Overall, there are several factors that might influence the number of transfers inferred from a set of gene families. Errors in the inferred gene or species trees will tend to inflate the number of inferred transfers, because disagreement between gene and species trees is typically more parsimoniously explained by transfers than by series of duplications and losses. As noted previously, processes such as hybridization (Szöllősi et al. 2015) and ILS (Bremer et al. 2022; Morel et al. 2022) will also produce disagreements between the gene and species trees that might be taken as evidence of gene transfer, although the ALE, GeneRax and SpeciesRax models have been show to be to be relatively robust to ILS (Morel et al. 2022). Likewise, previous work has indicated that reconciliation methods tend to infer fewer spurious transfers (and indeed, other kinds of events) than species tree-unaware methods, because reconciled gene trees are more accurate than gene trees inferred from the multiple sequence alignment alone; see (Szöllõsi et al. 2013; Scornavacca et al. 2015; Morel et al. 2020) for simulations quantifying the increased accuracy of reconciled gene trees, which appears to arise because information from the species tree can help to correctly resolve ambiguous regions of single gene trees. When comparing reconciled and species tree-unaware gene trees, we observed 24%, 59%, and 46% reductions in the mean numbers of duplications, transfers, and losses per gene family in an empirical dataset comprising 36 cyanobacterial genomes (Szöllõsi et al. 2013; Morel et al. 2020).

Another factor that is likely to influence the relative number of inferred transfers and duplications is the density of taxon sampling. As pointed out by Tria and Martin (2021), we would expect denser taxon sampling to result in a higher proportion of inferred transfers; this is because, as taxa are sampled more closely, some apparent duplications are revealed to actually be transfers from close relatives. We note that any method that uses phylogenetic trees to distinguish short-distance transfers from duplications (whether by reconciling the gene tree against a species tree, or by parsing gene trees for incongruent relationships) can only do so when taxon sampling is dense enough to include relatives of the donor, recipient, and enough intermediate taxa to shift the balance of evidence from duplication-followed-by-loss to transfer. By parsing gene trees inferred under site-homogeneous substitution models from a large sample (5,655) of prokaryotic genomes, Tria and Martin inferred an overall 50-fold excess of transfers over duplications. It will be interesting to compare reconciliation-based and species tree-unaware estimates of gene transfer on these larger datasets. We predict that numbers of inferred short range transfers will be substantially higher than the values plotted in Figure 3 on more densely-sampled bacterial datasets under reconciliation analyses, although performing these analyses with the best-fitting substitution models is currently challenging due to issues of computational tractability.

In sum, the conclusion that ALE recovers similar frequencies of transfer and duplication in Bacteria and eukaryotes (Bremer et al. 2022) is incorrect, and may have been due to misinterpretation of the ALE output, as illustrated in Figure 3 and discussed in more detail in Supplementary Text. Further, we note that the suggestion that inferred duplication, transfer and loss rates are unrealistic because they imply an excess of gene gains over losses through time (Bremer et al. (2022)) is also incorrect. We summarise and discuss reconciliation-based inferences of gene gain and loss on the bacterial dataset in the Supplementary Text.

### Constraining the ratio of duplication and transfer parameters to a predefined value reduces model fit and performance, particularly for the most informative gene families

Frequencies of gene duplication, transfer and loss vary across gene families. In ALE, the default procedure is to model these processes with three separate parameters for each gene family. Bremer et al. (2022) investigated the impact of constraining the ς and τ parameters on root inference. First they estimated the parameters of the model using maximum likelihood, as in the original analyses (Coleman et al. 2021). Using ML, they recovered the same root region as Coleman et al. (2021). On their test dataset of opisthokonts (fungi and metazoa), the default approach recovered the expected root (between fungi and metazoa) with maximum support. These results support the notion that reconciliation models can recover accurate root information when the model parameters are estimated from the data. For clarity, we note that Bremer et al. (2022) incorrectly refer to maximum likelihood (ML) parameter estimates as “1:1 T:D ratio” throughout their study; however, the unconstrained ς, τ and λ parameters are freely estimated via ML and do not, in general, have a 1:1 ratio; see Figure 3(i).

Next, Bremer et al. performed an experiment in which they fixed the ratio of τ and ς parameters. They did not, however, perform ML under a constrained ratio. Instead, they first estimated all three parameters freely by ML, and then performed a second analysis fixing ς as a multiple of the estimated τ (for example, ς =0.02*τ for the 50:1 τ: ς case). For the bacterial dataset, these analyses resulted in a loss of power, with the set of root positions that could not be rejected expanding to include additional nearby branches of the species tree. The same effect was observed on the opisthokont dataset for the 1:2 and 1:50 τ: ς cases. Interestingly, when the rates were fixed to a highly implausible 50-to-1 ratio of τ to ς in opisthokonts, the true root was no longer recovered in the credible set.

In their experiments, Bremer et al. (2022) did not investigate the impact of constraining the ς and τ parameters on model fit. Statistical analysis is usually best done under the best-fitting model (Sokal & Rohlf), and when the model parameters were estimated using maximum likelihood, ALE recovered the generally accepted fungal-metazoan root in the opisthokont data (Bremer et al. 2022). Therefore, it seems possible that the loss of precision and accuracy observed when constraining the ς and τ parameters might result from a loss of model fit.

To investigate the impact of fixing parameter ratios on model fit, we first implemented the ability to fix DTL parameter ratios in ALE. Optimising τ, ς and λ allows for a valid statistical comparison that is fairer to the simpler model; we hope that this additional function in ALE will also be of use for future investigations of genome evolution. This “fixed TD” model, which fixes the ratio of τ, ς to a user specified value, infers one fewer parameter per gene family than the full model used in the original analyses.

Having implemented the model proposed by Bremer et al., we then compared gene family likelihoods for the 11272 gene families under the full (independent ς, τ, λ parameters) and restricted model. Model fit was substantially worse under the restricted model; in the opisthokont dataset, fixing ς = 2τ resulted in a mean reduction in log likelihood of 14.6 units per gene family, while in Bacteria fixing ς = (1/50) (the “50:1” ratio) resulted in a mean loss of 20.2 log likelihood units per family; Table 2. Joint estimation of the single ς parameter (with τ set according to the prescribed ratio) and λ by ML using the new implementation in ALE greatly improved the fit of the simple model compared to Bremer et al.’s approximation, although model fit was still significantly worse than the default approach in which ς, τ and λ are all estimated independently (Tables 1 and 2). These results suggest that the loss of power reported by Bremer et al. is due to the use of an overly-simple model that fits the real data substantially worse than the default approach.

**Table 1:** Fixing τ:ς across gene families results in a significant loss of model fit in the fungi-metazoa dataset. Due to computational limitations, these values are computed for 15,556 of the original 15,614 families, because the model could not be fit for 58 families under the 1:50 T:D condition due to numerical instability; inferences are closely similar for the remaining families on the three other datasets. The first row shows the summed gene family likelihood when DTL parameters are independently estimated by maximum likelihood (the default setting). The second row shows the impact of setting ς to twice the value of the value of the parameter estimated by ML in the initial analysis (as per Bremer et al. 2022); this results in a large reduction in model fit. Subsequent rows show the log likelihood summed over familles when τ:ς was set to a fixed ratio, but the value of this joint τς parameter was estimated by ML. AIC was calculated as 2(total number of parameters)-2(summed log-likelihood); the default approach provides the best model fit (lowest AIC).. The final column summarises the number of families that reject the simpler model on a per-family basis).

| $\tau:\delta$ ratio | Summed LL | ΔLL | ΔLL/family | AIC | Families that reject simpler model (AIC) |
| --- | --- | --- | --- | --- | --- |
| Free (maximum likelihood DTL estimation) | -507684.73 |  |  | **1062037** |  |
| Estimate free, then fix $\delta=2*\tau_{\text{opt}}$ | -735742.10 | 228057.37 | 14.66 | 1502596 | 7335/15556 |
| 1:2 | -534420.18 | 26735.45 | 1.71 | 1099952 | 4817/15556 |
| 1:50 | -534009.37 | 26324.64 | 1.69 | 1099131 | 4309/15556 |
| 50:1 | -669876.28 | 162191.54 | 10.4 | 1370865 | 6895/15556 |

**Table 2:** Fixing τ:ς across gene families results in a significant loss of model fit in the bacteria dataset. The first row shows the summed gene family likelihood when DTL parameters are estimated by maximum likelihood (the default setting). The second row shows the impact of setting ς to one-fiftieth the value of the value of the τ parameter estimated by ML in the initial analysis (as per Bremer et al. 2022); this results in a large reduction in model fit. Subsequent rows show the log likelihood summed over familles when τ:ς was set to a fixed ratio, but the value of this joint τς parameter was estimated by ML. AIC was calculated as 2(total number of parameters)-2(summed log-likelihood); the default approach provides the best model fit (lowest AIC). The final column summarises the number of families that reject the simpler model on a per-family basis).

| $\tau:\delta$ ratio | Summed LL | ΔLL | ΔLL/family | AIC | Families that reject simpler model (AIC) |
| --- | --- | --- | --- | --- | --- |
| Free (maximum likelihood DTL estimation) | -2204764.16 |  |  | **4443344** |  |
| Estimate free, then fix $\delta=0.02*\tau_{\text{opt}}$ | -2432608.65 | 227844.49 | 20.21 | 4887761 | 5930/11272 |
| 50:1 | -2318604.50 | 113840.34 | 10.1 | 4659753 | 5908/11272 |
| 100:1 | -2319889.26 | 115125.09 | 10.21 | 4662323 | 5687/11272 |

To systematically assess model fit on a per-family basis, we used the Akaike Information Criterion (AIC) to compare support for the simpler (2-parameter) versus the more complex and hence parameter rich (3-parameter) model for each gene family under each condition. This analysis indicated that the AIC rejected the simpler model for 28-52% of gene families across the range of ratios tested (Tables 1 and 2), when considered individually. One contributor to the preference of individual families for the simple or more complex model is family size: in the bacterial dataset, families for which the AIC rejected the simple model tended to be larger (median 11 and mean 47.06 gene copies, compared to median 8 and mean 24.1 for families that did not reject the simple model by AIC; *p* = 4.84 · 10^−57^, Wilcoxon rank-sum test), and family size was strongly correlated with the strength (log likelihood difference between the simple and complex model) with which the simpler model was rejected (Spearman’s ρ = 0.23, *p* < 2.2 · 10^−16^). This result suggests that independent estimation of ς, τ, and λ parameters is particularly important for larger gene families, while for the smallest gene families the amount of data does not appear to suffice to reliably optimise them.

To assess whether the larger gene families for which AIC individually rejects the simple (fixed τ: ς ratio) model have distinct rooting information from the smaller families that do not, we divided the families of the bacterial dataset into these two sets and performed an AU root test separately on each. In the original analysis (Coleman et al. (2021)), we obtained support for a root region including three branches, corresponding to a root between Gracilicutes and Terrabacteria or on the adjacent branch leading to Fusobacteriota; the analysis did not distinguish whether Fusobacteriota branched as sister to Gracilicutes or to Terrabacteria. The AU test on the 5930 (fixed two-step procedure) or 5908 families for which AIC rejected the simple model recovered a root region similar to that inferred from the full dataset, with a root either between Gracilicutes and Terrabacteria or on Fusobacteriota (Figure SX). Interestingly, this root region contained one fewer branch than the test on the full data, with the branching of Fusobacteriota on the terrabacterial side of the root rejected at *P* < 0.05. That is, the analysis placed additional weight on Fusobacteriota as the earliest-branching group within Gracilicutes, a position that is consistent with analyses of some cell envelope characters (Fusobacteriota possess a Gracilicute-type system for tethering the outer membrane to the cell, (Witwinowski et al. 2022)). By contrast, the AU test on the 5342 smaller gene families for which AIC did not reject the simpler model was much less informative, with a root region including the Gracilicute-Terrabacteria divide (with Fusobacteria branching at the root of Terrabacteria) but also 10 other positions (Figure S2). In sum, these analyses suggest that reconciliation model fit for larger gene families is optimal using the default approach in which ς, τ, and λ are optimised independently, and that such families also contain much of the rooting signal that is available to reconciliation analyses.

The real evolutionary process is more complex than the best available models, and so parameter inferences and analyses under even the full D, T, L model are, to some extent, misspecified. In this context, the experiments of Bremer et al. (2022) on the opisthokont dataset are encouraging. When parameters were estimated from the data, the most plausible root was recovered with maximum support. The main effect of model misspecification appears to be a loss of statistical power, with the model being unable to differentiate between additional branches as fit worsened (Table 1). Only when the TD parameters were set to very implausible values (a 50-fold higher than ς in animals and fungi, for all gene families) did the analysis become misleading, in the sense that the expected root was no longer in the 95% credible set. These analyses suggest that the best approach for empirical analyses is to estimate model parameters from the data, rather than setting them to subjective values.

### The nature of the root signal in reconciliation analyses

Different root positions on the species tree imply different probabilities for different reconciliation scenarios, and this property enables the root of the species tree to be inferred using reconciliation methods. It is therefore interesting to consider the nature of the root signal that is being captured in reconciliation analyses. Under the assumption that it is caused by DTL events that are captured by the model, this signal is derived from varying degrees of incongruence with different rooted species trees. As a result, gene families that experience very few DTL events have very limited power to identify the root of the species tree, because a perfectly congruent gene tree is equally consistent with all species tree root positions. This explains why the analysis of the single-copy marker genes from the fungi-metazoa dataset did not distinguish the root of the tree in Bremer et al. (2022)’s experiments; these genes have experienced no inferred duplications and very few inferred transfers or losses (Table 1).

The classic case in which the tree of life was rooted using pre-LUCA duplications (Iwabe et al. 1989; Gogarten et al. 1989; Brown & Doolittle 1995) demonstrates how gene duplications inform root inference, but transfers and losses can also provide information on the root. Indeed, the information provided by all three kinds of events is similar: a duplication, acquisition by transfer, or loss at the base of a clade on the species tree provides evidence that the root of the tree is not within that clade (Iwabe et al. 1989; Gogarten et al. 1989; Lake et al. 2009; Szöllosi et al. 2012; Williams et al. 2017). To investigate how inferred DTL events and different kinds of gene families contribute to root signal in a given empirical dataset, several approaches seem possible. In Coleman et al. (2021), we explored the nature of the root signal by ranking gene families by a range of different metrics, then sequentially removing families and evaluating the impact on root support. These analyses indicate that broadly-distributed and predominantly, but not entirely, vertically-evolving gene families were the most informative, because filtering these families from the dataset reduced the difference between the log-likelihood scores of the different root positions. For example, filtering out the top 20% of mcl gene families ranked by verticality or breadth of distribution in extant Bacteria (number of genomes encoding the gene family) greatly reduced the likelihood difference among candidate root positions, while removing the bottom 20% of gene families by this criterion had no effect; see Figure S12 in Coleman et al. (2021). Since such families are expected to have originated early along the species tree, this finding is consistent with gene tree-species tree incongruence caused by deep DTL events driving the root signal.

### Inferring ancestral gene content using reconciliation methods

The outcome of a gene tree-species tree reconciliation analysis is a mapping of gene family evolution onto the branches of the species tree. As such, the results can be used directly for obtaining probabilities for the presence of a set of gene families at internal nodes of interest on the species tree. Specifically, with probabilistic reconciliation methods such as ALE and GeneRax, the proportion of sampled reconciliations in which a gene family was present at a given node (the presence probability, available in the ALE reconciliation file output) is a natural metric for performing ancestral gene content analyses; for example, if a gene family was present at the root of the tree in 90 out of 100 sampled reconciliations, the root presence probability (PP) is 0.9 (note that reconciliation scenarios and model parameters are estimated jointly, so reconciliations are sampled proportionally to their probability under the ALE model). The number of genes present at an internal node of the tree can be estimated by summing the estimated copy numbers over all gene families (summing partial copies averages over the uncertainty in the identity of the families present). Since some gene families present at the internal nodes of the tree have since gone extinct, basing gene repertoire size directly on these counts will result in an underestimate. One possibility is to estimate the probability that a gene family at a given node has since gone extinct, based on the inferred model parameters; the gene content estimate from surviving families can then be corrected accordingly.

To obtain the set of gene families estimated to be present at an ancestral node, a reasonable PP threshold — such as 0.5 or 0.95 — can be used. This approach to reconciliation-based ancestral reconstruction has been used in a number of published analyses to investigate gene content evolution on the internal nodes of a species tree (Williams et al. 2017; Martijn et al. 2020; Coleman et al. 2021; Harris et al.; Dharamshi et al. 2023). Dharamshi et al. (2023) developed a snakemake-based workflow for performing these analyses using ALE that may be of interest (https://github.com/maxemil/ALE-pipeline).

An important feature of reconciliation-based methods for ancestral gene content reconstruction is that they consider the gene family phylogeny in the analysis. This is distinct from some alternative methods for studying gene content evolution, such as Count (Csurös 2010), CAFE (Mendes et al. 2020) and GLOOME (Cohen et al. 2010), which model the evolution of phylogenetic profiles (the number of gene copies in genomes) on a rooted species tree. Like ALE and GeneRax, these methods use probabilistic models to describe the evolution of gene families (in this case, gene family profiles) along the species tree as a series of duplication, transfer, loss and origination events. One benefit is that this formulation allows rate variation across the branches of the species tree to be modelled, a current shortcoming of ALE/GeneRax (see Conclusion). However, incorporating phylogenetic information has a profound effect on inferences of ancestral gene content (Figure 4). By basing inferences on the reconciliation of the gene and species trees, ALE and GeneRax are better able to detect gene transfers in deep time (on the basis of gene tree-species tree incongruence), and as a result are conservative compared to profile-based approaches, which are only able to detect transfer events that leave a clear signal in the number of gene copies at the tips of the species tree. This, however, also implies an important caveat regarding reconciliation-based gene contents: reconstruction errors or phylogenetic biases in gene trees will be interpreted as spurious DTL events, and as illustrated in Fig 4c result in an systematic underestimation of the number of gene copies at deeper nodes of the species tree.

**Figure 4:**
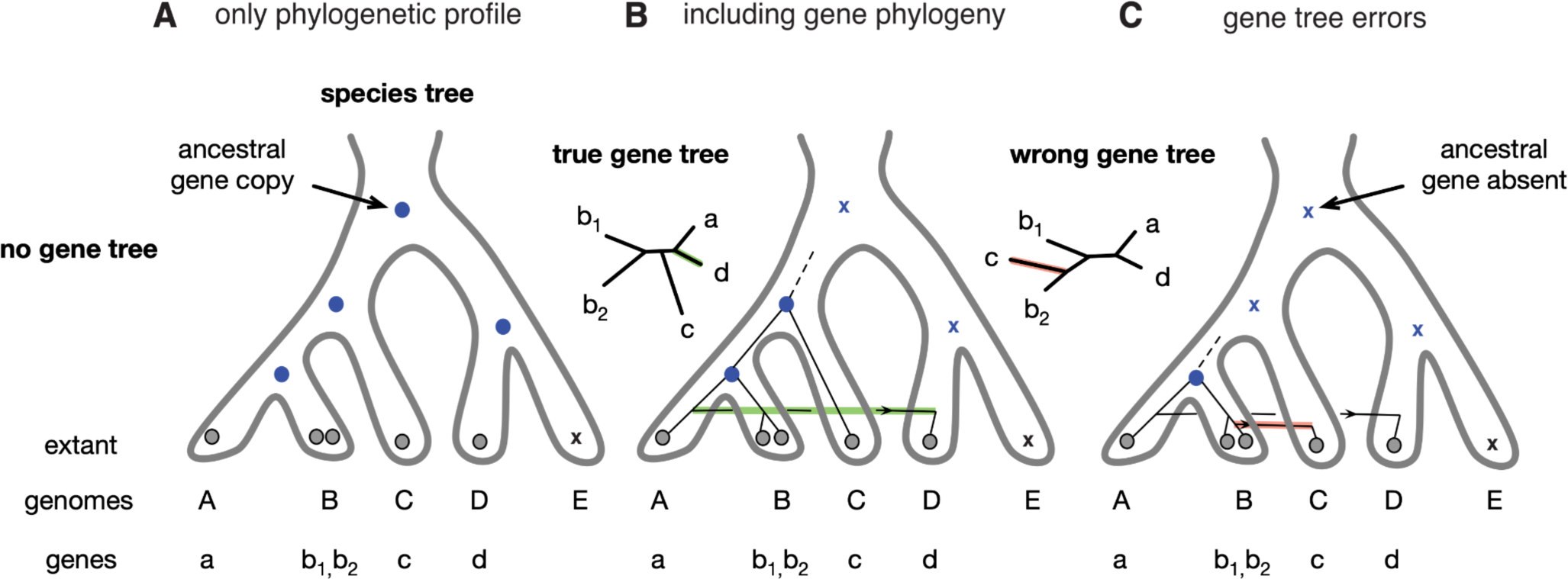
The effect of incorporating gene tree topology information when inferring ancestral genome contents. Grey dots denote observed gene copies in extant species, blue dots denote inferred ancestral presence of a gene, blue Xs denote inferred ancestral absence of a gene. Consider a gene family broadly distributed in extant taxa (A). On the basis of this phylogenetic profile, it appears likely that the gene traces to the root of the species tree. However, comparison of the gene tree to the species tree (B) suggests a recent horizontal acquisition of the gene on the right hand side of the species tree root; as a result, the gene is inferred to have originated more recently. Comparison of (A) and (B) illustrates why profile methods may tend to overestimate ancestral gene contents compared to reconciliation-based methods. (C) Errors in the reconstructed gene tree may be spuriously interpreted as additional gene transfers, so that the evolutionary age of the gene is under-estimated. This case illustrates why reconciliation-based methods may tend to under-estimate ancestral gene repertoires. This figure is based on that of (Kellner et al. 2018) with some modifications.

Although ALE and GeneRax aim to minimise the effect of stochastic gene tree errors by inferring gene trees combining phylogenetic information from both the gene family multiple sequence alignment and the species tree (cf. e.g. Fig. 2a-c in (Szöllõsi et al. 2013)), reconciliation methods remain sensitive to phylogenetic errors resulting in systematic biases, such as inadequate modelling of the substitution process, as well as upstream errors in the inference of gene families that can result in missing genes or spurious homologs. As a result, the systematic underestimation of gene copies at deeper nodes of the species tree remains a substantial challenge, particularly at very deep phylogenetic scales.

One solution to this issue is to pool phylogenetic signal across genes, either by concatenation or by jointly estimating presence probabilities for sets of genes. For example, Coleman et al. (2021) estimated a single root origination probability for each COG category of genes, then calculated individual root probabilities for each gene based on its reconciliation with the species tree and the root origination probability for its class. In practical terms, the approach appears promising, resulting in a last bacterial common ancestor with a gene repertoire within the range of extant Bacteria, high probabilities for core cellular machinery such as the transcription and translation systems, and conserved metabolic pathways such as glycolysis, the TCA cycle and the pentose phosphate pathway. On the other hand, this kind of approach is clearly sensitive to the way that signal is pooled across genes and as a result, less conservative than the standard reconciliation approach.

### Conclusions: limitations of current reconciliation methods, and prospects for progress

Reconciliation methods offer a powerful approach for bringing together phylogenomics and comparative genomics, studying the processes of evolution, and addressing major unanswered questions about the evolution of cellular life as far back as LUCA. But the existing models are still relatively crude, and we would like to conclude by highlighting current limitations and some ways in which these might be overcome. An open question is how best to model the ways in which frequencies of gene duplication, transfer and loss vary across the tree of life. DTL frequencies vary enormously across families (Jain et al. 1999; Cohen et al. 2011) and so, as shown above, a simple model in which parameters are fixed to a global ratio fits real data poorly. Estimating the ς, τ and λ parameters separately and independently for each gene family provides a better fit, but might induce over-parameterisation, especially for smaller families. There is a clear parallel here with the problem of estimating per-site evolutionary rates in traditional phylogenetics, where mixture models have been used to capture rate variation across sites (potentially, among families) without the need to estimate large numbers of parameters; for example, on the assumption that rates are gamma-distributed, only a single parameter (the alpha shape parameter) is needed. More flexibly, and as in the “free rates” model (Minh et al. 2020), gene family rates could be modelled as a mixture of rates, with the number of rate categories chosen on the basis of model fit (for example, using the Akaike or Bayesian Information Criteria (Posada & Buckley 2004)). One current difficulty in conducting inferences under this kind of model is that, in comparison to estimating site rates, the problem is three-dimensional (i.e. one dimension for each of the ς, τ and λ parameters, and reconciliations for all gene families would need to be recomputed each time one of the shape parameters was updated). An approximate solution might be to divide gene families into categories corresponding to similar ς, τ and λ values, which could then be fixed for all families within a bin.

An additional limitation of the current ALE/GeneRax model is the simplifying assumption that the same ς, τ and λ parameters apply to all branches of the species tree. This assumption is certainly violated by real data: for example, vertically inherited endosymbionts and intracellular parasites often undergo extensive gene loss compared to their free-living relatives (McCutcheon & Moran 2011), while multicellular eukaryotes are commonly assumed to acquire fewer genes by horizontal transfer than do their unicellular relatives. Since inferred transfers, duplications and losses ultimately depend on the gene tree topologies, reconciliation analyses can recover these broad patterns in the variation of D, T and L across clades (for example, the higher T/D in Bacteria than Opisthokonts, and in Fungi compared to Metazoa - Figure 2 this study; but also (Ocaña-Pallarès et al. 2022)). However, the assumption of a constant branch-wise probability means that the method lacks the power to identify precisely where major shifts in the frequency of duplications, transfers or losses occur. In ALE, it is currently possible to test hypotheses about branch-wise shifts in D, T or L parameters by applying multipliers to specific branches of interest, and current work is focused on implementing “highways” of transfer between distant points on the species tree. However, a more general solution, involving optimization of parameters across branches and gene families, remains intractable. A first step in this direction would be to introduce a mixture model with a few branch specific categories.

Comparison and critique of current methods will be necessary for progress and to guide future method development. Despite their shortcomings, existing methods do capture broad-scale patterns in rates of duplication, transfer and loss across clades, and there is promising agreement between reconciliation methods and alternative rooting approaches in cases where there is biological consensus on the root (such as the root of plants or opisthokonts). An interesting open question is the extent to which frequencies of gene transfer vary across the tree of life. Within eukaryotes, published analyses suggest that transfers are less frequent in multicellular groups than in unicellular groups. This is clearly the case among animals, perhaps because of the germ/soma distinction (Ocaña-Pallarès et al. 2022) and also appears to hold for plants, where Harris et al. (2022) found transfers to be significantly less frequent than duplications in 31 high-quality streptophyte genomes (e.g., median branchwise transfers/duplications 0.21 for multicellular plants, median T/D 1.11 for their closest algal relatives; Harris et al. (2022)). By contrast, gene transfers appear relatively common in Fungi, the eukaryotic lineage in which gene transfers have been most extensively studied (Richards et al. 2011; Szöllősi et al. 2015), but frequencies may be still higher in other lineages such as Rhizaria, where a recent phylogenetic study suggested that 30% of gene families might have been acquired horizontally from prokaryotes or from other eukaryotes (van Hooff & Eme 2023). Reconciliation methods are useful for studying how these frequencies vary across clades (Szöllősi et al. 2015; Ocaña-Pallarès et al. 2022) and testing hypotheses about the underlying biology. It seems clear that as methodology improves, probabilistic reconciliation is poised to play an increasingly important role in our understanding of biological evolution.

## Data availability

Supplementary data for this manuscript are available in the Zenodo.org repository at DOI: 10.5281/zenodo.7682207.

## Acknowledgements

This work was supported by the Gordon and Betty Moore Foundation through grant GBMF9741 to TAW, AS, LS, and GJSz, and by a Royal Society University Research Fellowship to TAW. Furthermore, GJSz and LS received funding from the European Union’s Horizon 2020 research and innovation programme (grant agreement No. 714774, GENECLOCKS), A.S. has received funding from the Swedish Research Council (VR starting grant 2016-03559), the NWO-I foundation of the Netherlands Organisation for Scientific Research (WISE fellowship) and the European Research Council (ERC) under the European Union’s Horizon 2020 research and innovation programme (grant agreement No. 947317, ASymbEL). AAD and PH are supported by an Australian Research Council Laureate Fellowship (grant FL150100038). This work was financially supported by the Klaus Tschira Foundation, by DFG grant STA 860/6-2, and by the European Union’s Horizon Europe ERA Chair program under grant agreement No 101087081 (Comp-Biodiv-GR).

