## Supplementary Figures for "The power and limitations of species tree-aware phylogenetics"

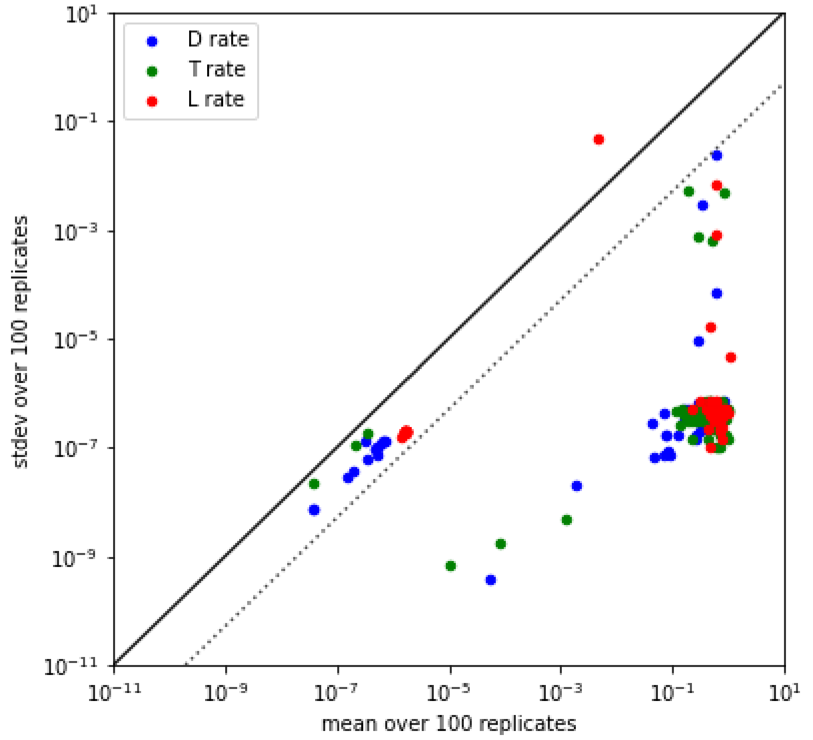


**Supplementary Figure 1: Maximum-likelihood estimation of duplication ($\delta$), transfer ($\tau$), and loss ($\lambda$) parameters in ALE is robust to the starting values used in the calculation.** We sampled 100 gene families randomly from the dataset of Coleman et al. (2021), then estimated $\delta$, $\tau$ and $\lambda$ parameters 100 times for each family, starting the ML optimization from random initial seeds each time. The plot shows the mean (x-axis) and standard deviation (y-axis) of the parameter estimates. The solid line corresponds to standard deviation = mean, while the dashed line denotes standard deviation = 1% of the mean. The results show that $\delta$, $\tau$ and $\lambda$ parameter estimates are robust to the starting seed values, with standard deviation < mean (typically, standard deviation << mean) in all but a single case (discussed below). The mean standard deviation of the gene family likelihoods across these 100 families was 0.00014 (median 0.0) log likelihood points. For the single outlier (the loss rate estimated for one family), the mean L rate is 0.0046 and the standard deviation is 0.046; for 99/100 replicates, L ~ 0 ($1 \times 10^{-10}$) with log-likelihood -18.91, whereas in one replicate L ~ 0.46 and log-likelihood -18.77, suggesting the optimization algorithm failed to find the maximum likelihood parameter configuration in this single case.

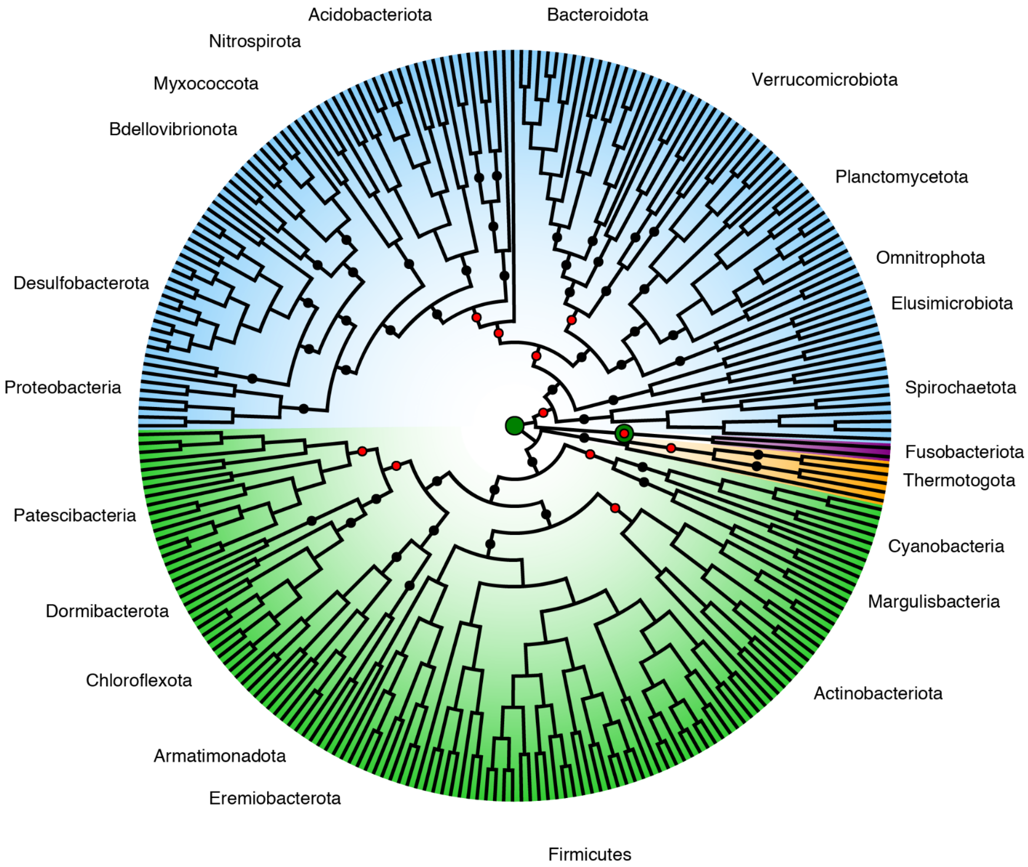


**Supplementary Figure 2: Rooting analysis using different sets of parameters.** We divided the gene families used in Coleman et al. (2021) into two groups: those that preferred a model with 3 parameters and those that preferred a model with 2 parameters in which the $\tau$ and $\delta$ parameters are fixed to a constant ratio. The 5930 families that prefer the 3 parameters support a root region indicated by the large two green dots. The 5342 that preferred the 2 parameters model (represented by the red dots) are in general smaller families and the root region supported by them is more diffuse. The remaining tested roots that were rejected by both sets of families appear as black dots. In this analysis, we divided genes into the two sets using the two-step inference procedure of Bremer et al. (2022), but dividing families according to their preference after jointly optimising $\tau:\delta$ (with 5908 families rejecting the simple model by AIC) produced the same result.

### Supplementary Text

*Genome evolutionary dynamics are best captured by branch-wise metrics*

In the following, we provide a more detailed explanation of the distinction between $\delta$, $\tau$ and $\lambda$ parameters (estimated per-family in ALE/GeneRax); the number of inferred duplication, transfer and loss events per-family (which result from analysing the data using the model); and the number of inferred duplication, transfer and loss events per branch of the species tree, to supplement the discussion of Figure 2 in the main text.

Frequencies of duplication, transfer and loss vary across gene families (e.g., ribosomal proteins are transferred less frequently than metabolic genes (Jain, Rivera, and Lake 1999)) and across lineages (gene transfers appear to be more frequent in Bacteria than in eukaryotes). The $\delta$, $\tau$ and $\lambda$ parameters in ALE are estimated for each family, but do not vary over the species tree. Starting from gene trees, doing inference with this model gives rise to inferred duplication, transfer and loss events that map to branches of the species tree. These lineage-specific events can then be summarised to give branch-wise counts of inferred duplication, transfer and loss events that can be used to compare genome dynamics between lineages. The $\delta$, $\tau$ and $\lambda$ parameters, however, are not representative of lineage-level dynamics (or across-lineage variation) because they are estimated family-wise (Figure 3). Each family-wise ALE output file lists inferred events for each branch of the species tree, and branch-wise counts of inferred duplications, transfers and losses are a natural way to summarise these inferences; these various parameters and summary statistics are plotted in Figure 3; a Python script to perform these calculations is provided at https://github.com/AADavin/ALEtutorial. In their critique, Bremer et al. (2022) apparently did not inspect per-genome inferences using ALE; instead, it appears that they compared family-wise $\delta$ and $\tau$ parameters to per-genome counts of events from previous analyses.

A second potential source of error relates to using overall averages to summarise gene family dynamics, when in fact gene family size and $\delta$, $\tau$ and $\lambda$ parameter values vary greatly across families. For example, the global means of $\tau$ and $\delta$ across all families in the bacterial dataset are 0.42 and 0.19, respectively, and this might indeed suggest that frequencies of transfer and duplication are roughly 2:1 in this dataset. However, the global means (or medians) of these parameters do not capture the substantial variation in frequencies of duplication, transfer and loss across families: note that the means of the $\tau/\delta$ ratios for Bacteria and opisthokonts are both very far from the ratio of the means (Figure 2), and it is these per-family dynamics that underlie inferred events. We also note that Bremer et al. seem to have calculated metrics based on model parameters, not on model inferences — that is, inferred events. We would recommend metrics based on inferred events, rather than model parameters. The fundamental reason is that inferences have a clear biological interpretation, whereas model parameters are of secondary interest and, in a data-homogeneous model, do not reflect biologically interesting variation in the process we are attempting to model. Consider an analogy to studying substitution histories in standard phylogenetics: we might wish to know which substitutions occurred where (on which branch of the tree), or how many substitutions occurred per year or along a branch of interest. In a homogeneous substitution model, the equilibrium frequencies of the amino acids and the instantaneous rates of change among them are fixed. Despite this, inference using the model reveals variation in branch lengths, rates and patterns of substitution across the tree. These patterns reflect information in the data that is captured by the model. In the same way, the most relevant output from fitting a reconciliation model is the evolutionary scenario and the associated inferred duplication, transfer and loss events which, like substitutions in the standard phylogenetic analysis, occur along branches of the species tree.

Beyond the parameter/inference distinction, there is an additional drawback to metrics based on per-family ratios: many families experience 0 duplications (e.g., 5,314/11,272 families in the bacterial dataset), so their T/D (expressed as inferred events) is infinite, and $\tau/\delta$ tends to infinity ($\delta$ ~ 0 when zero duplications are inferred); similarly, families with 0 transfers (of which there are 9673 in the fungal-metazoan dataset) will have $\tau/\delta$ ~ 0. Thus, family-wise T/D and $\tau/\delta$ range from 0 to infinity. This does not present a problem for inference using ALE, because the per-family $\tau$ and $\delta$ parameters are estimated separately; however, it makes the per-family parameter ratios a noisy, unstable and potentially misleading metric. For example, the median per-family ratio of $\tau$ to $\delta$ is 0 for the Fungi-Metazoa dataset and 7 for the Bacteria dataset, while the means are $7\text{x}10^7$ for Fungi-Metazoa and $1\text{x}10^9$ for Bacteria; the enormous difference between means and medians demonstrates that these summaries inadequately describe the evolutionary dynamics of the underlying gene families. Bremer et al.'s remark (Bremer et al. 2022) that the range of per-family T/D rate ratios inferred by ALE is unreasonably wide is most likely based upon conflating per-family $\tau/\delta$ ratios (which they calculated for the ALE analyses) with per-*genome* counts of inferred *events* — transfers and duplications — to which they compared ALE's per-family values. Perhaps the clearest example of why expressing evolutionary dynamics as a $\tau/\delta$ ratio can be unhelpful comes from the analysis of the 117 single-copy marker genes of the opisthokont dataset. Bremer et al. (2022) reported that the mean $\tau/\delta$ for these genes was 26968:1, which they regarded as abnormally high. However, the high ratio simply results from the fact that ALE infers 0 duplications for all 117 families; ALE estimates verticality at 0.99 for these genes, which seems reasonable for a set of generally vertically-evolving marker genes (Figure 3).

This conflation of family-wise $\tau$ and $\delta$ parameters with branch-wise T and D events also appears to underlie some previous critiques of ALE by some of the same authors. For example, Tria and Martin (2021) claimed that ALE inferred 1-2 transfers per gene duplication in a previous analysis of Cyanobacteria (Szöllősi et al. 2015). In fact, branch-wise T/D was 3.37 (median) and 6.6 (mean), as plotted in the original study (Figure 4, Szöllősi et al. 2015, where ~79% of inferred events were transfers).

*Reconciliation-based estimates of genome size evolution*

In a further critique of reconciliation-based analyses, Bremer et al. investigated the per-family ratio of the $\lambda$ to gene gain rate (which they defined as $\tau+\delta$) in the 11,272 families of the bacterial dataset. A “loss/gain ratio” > 1 denotes a gene family with a general tendency to contract over time, while a ratio <1 indicates a family that tends to expand over time (that is, where family expansion by gene duplication, or acquisition by transfer, outweighs losses). They reported a mean loss/gain ratio of 0.76, which they argued was unrealistic given that bacterial genes are lost more often than they are acquired (Bremer et al.). This calculation and inference appear to be incorrect, as we briefly explain below.

Bremer et al. (2022) calculated the loss/gain ratio by dividing the mean $\lambda$ by the sum of the mean of $\delta$ and the mean of $\tau$, to obtain an estimate of 0.76. However, the ratio of two means is not the mean of their ratios, and in fact the actual mean $\lambda/\delta+\tau$ for the bacterial dataset ranges from $1.23\text{x}10^7$ - $1.37\text{x}10^7$ across the three branches of the root region; the median is 1 for all three root branches. The enormous difference between the means and medians arises because the data are not normally distributed: as discussed above, many families experience 0 Ds, Ts or Ls, and so family-wise ratios of parameters or events are highly unstable. Thus, judged by this $\lambda/\delta+\tau$ metric, losses generally equal or exceed gains among the gene families of the bacterial dataset.

However, it is important to note that this per-family $\lambda/\delta+\tau$ metric does not provide insight into changes in gene content over time, because it refers to per-family (not per-genome) $\delta,\tau$ and $\lambda$ parameters (not events). In order to track changes in gene content, we need to map gain and loss events onto the species tree. We also need to consider gene originations, an important source of new genes that is not considered in the $\lambda/\delta+\tau$ metric.

To summarise ALE-based estimates of gene content change through time, we calculated the branch-wise number of gene losses, divided by the sum of gene originations, duplications and transfers (L/O+D+T). The mean and median branch-wise values are 0.9213 and 0.8777, respectively, suggesting that gains slightly outweigh losses overall in the bacterial dataset (that is, on average ~0.92 losses occur for every gene gain resulting from an origination, acquisition by transfer, or gene duplication). Does this result imply that bacterial gene repertoires have increased continuously through time? No, because these values have been inferred from gene families that have survived to be sampled in the present day. Note that, for a gene family to go extinct, within-family losses must equal or exceed gains. Gene families with L > O+D+T are the only ones that go extinct, and so our present-day sample of families is biased towards families with lower L/O+D+T values; in other words, we would expect these values, given that we base our inferences on extant gene families. This effect can be readily seen in the bacterial dataset by plotting L/O+D+T as a function of their distance from the root of the tree (Figure S3): the ratio is lowest near the root, and rises above 1 on recent branches. This is because the families which inform L/O+D+T on a branch must have survived from that branch to the present day; the ratio therefore appears lower on more ancient branches.

This is not to say that current reconciliation models fully capture changes in genome dynamics. For example, in the ALE/GeneRax model, $\delta,\tau$ and $\lambda$ parameters are homogeneous over the tree as well as independent of gene family copy number. In reality, not only do rates of duplication, transfer and loss vary across branches, but there is strong evidence that they also depend on the number of homologous gene copies in a particular genome (Huynen and van Nimwegen 1998). Nonetheless, published ALE results seem plausible, identifying the large loss of gene content in the common ancestor of the CPR clade ((Brown et al. 2015; Méheust et al. 2019) and major gene losses near the tips of the tree leading to parasitic and endosymbiotic lineages, in agreement with alternative (non-reconciliation) approaches (McCutcheon and Moran 2011).

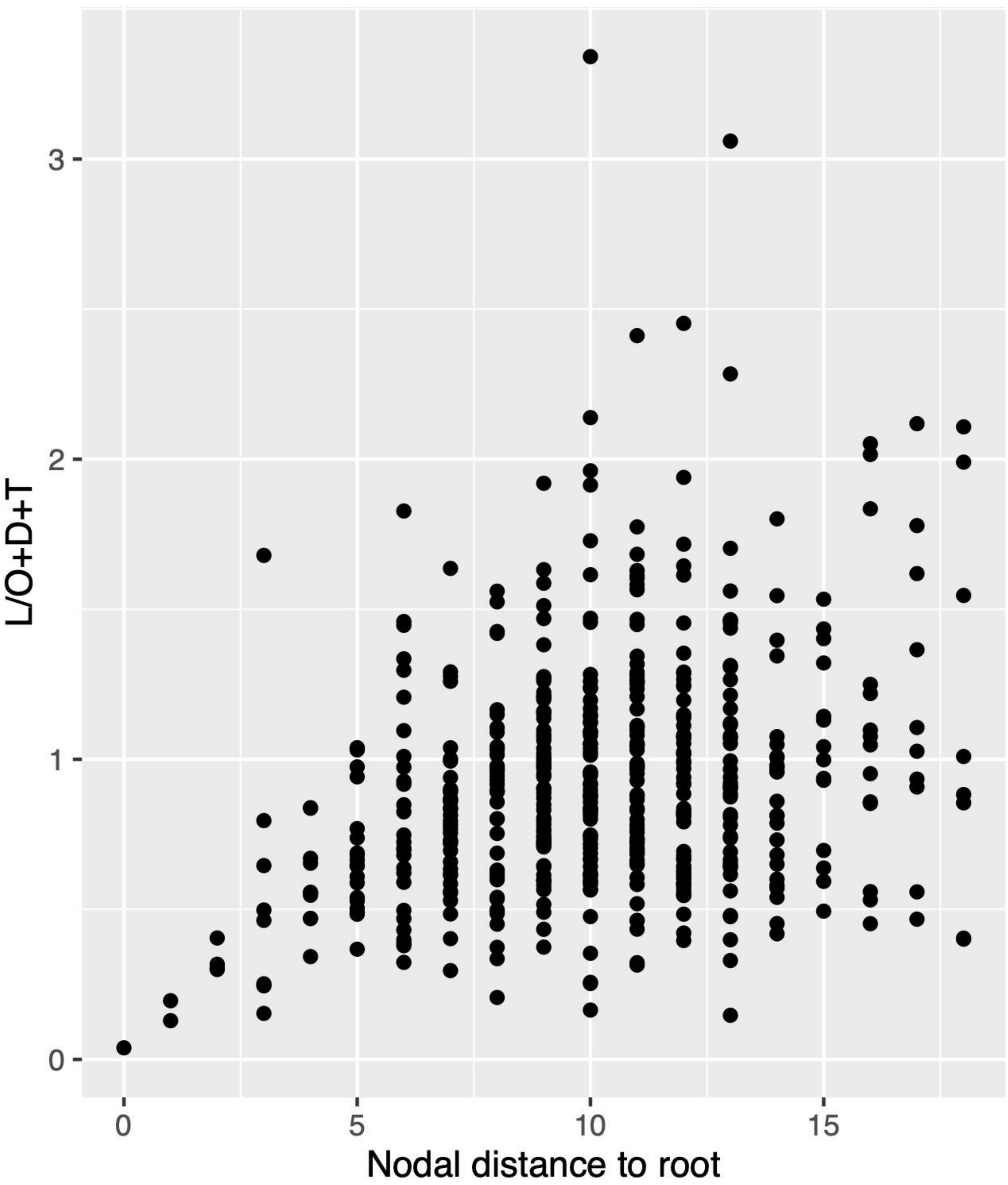


**Supplementary Figure 3: Survivorship bias in estimates of gene loss and gain in the bacterial dataset.** When calculated from gene families that have survived to the present day, the ratio of gene losses to gains (originations, duplications, and acquisition by transfer) along the species tree shows a clear survivorship bias: branches closest to the root experience more gains than losses, while the balance shifts on recent branches. This is because gene families present in the past that survived to the present day tend to experience more gains than losses; gene families in which losses exceed gains go extinct and are not sampled by present-day phylogeneticists.
